# Proprioceptive Genes as a Source of Genetic Variation Underlying Robustness for Flight Performance in *Drosophila*

**DOI:** 10.1101/2021.06.03.446923

**Authors:** Adam N. Spierer, David M. Rand

**Author notes:** **Corresponding authors**: David M. Rand, 80 Waterman St, Providence, RI 20912.

## Abstract

A central challenge of quantitative genetics is partitioning phenotypic variation into genetic and non-genetic components. These non-genetic components are usually interpreted as environmental effects; however, variation between genetically identical individuals in a common environment can still exhibit phenotypic variation. A trait’s resistance to variation is called robustness, though the genetics underlying it are poorly understood. Accordingly, we performed an association study on a previously studied, whole organism trait: robustness for flight performance. Using 197 of the Drosophila Genetic Reference Panel (DGRP) lines, we surveyed variation across single nucleotide polymorphisms, whole genes, and epistatic interactions to find genetic modifiers robustness for flight performance. There was an abundance of genes involved in the development of sensory organs and processing of external stimuli, supporting previous work that processing proprioceptive cues is important for affecting variation in flight performance. Additionally, we tested insertional mutants for their effect on robustness using candidate genes found to modify flight performance. These results suggest several genes involved in modulating a trait mean are also important for affecting trait variance, or robustness, as well.

**Article Summary:** We sought to understand the genetic architecture of robustness (variation in a trait caused by non-genetic factors) for flight performance. We used 197 Drosophila Genetic Reference Panel (DGRP) lines to find significant individual variants and pairs of epistatic interactions, many of which were involved in proprioception. Additionally, we validated significant genes identified from a prior study for the mean of flight performance, showing genes affecting trait means may also affect trait robustness.

## INTRODUCTION

Evolution acts on the genetic variation underlying phenotypic variation among individuals and populations. While many research programs focus on understanding genetic factors that contribute to phenotypic variation, fewer focus on non-genetic factors. The phenomenon of non-genetic (micro-environmental) variation describes the phenotypic variation that occurs in the absence of genetic variation, best studied in genetically identical individuals. Non-genetic variation can arise from external (environmental) or internal (developmental) factors. Phenotypic variation across different environmental conditions (e.g. temperature) in genetically homogenous organisms is termed phenotypic plasticity. However, significant phenotypic variation can also arise among genetically homogeneous organisms in the absence of explicit environmental variation (Morgante *et al*. 2015; Vogt 2015). Here, internal factors spur developmental noise in stochastic molecular processes, such as important transcripts or signals in very low abundance, which can result in varying levels of developmental stability (Albayrak *et al*. 2016; Schor *et al*. 2017; Klingenberg 2019). The processes or ability for organisms to maintain a consistent phenotype in the presence of these perturbations is termed buffering, while the resulting phenotype is deemed robustness (Klingenberg 2019).

Developmental noise can affect an organism’s developmental trajectory, which may impact the efficacy of natural selection by altering the association between genotype and phenotype. While it is difficult to directly observe developmental noise, deviations from an expected phenotype provide an adequate lens for study (Morgante *et al*. 2015; Vogt 2015). An example of this is deviations in bilateral symmetry (fluctuating asymmetry) (Valen 1962; Soto *et al*. 2008), which are hypothesized to be negatively associated with fitness in the case of facial symmetry (QUINTO-Sanchez *et al*. 2018; Lajus *et al*. 2019). Some genetic safeguards exist to buffer against developmental noise and maintain phenotypic robustness in the presence of these stressors. Chaperonins (HSP90) do so by maintaining a protein’s structure during stressful times (Rutherford and Lindquist 1998; Chen and Wagner 2012), as does the mitochondrial unfolded protein response in maintaining homeostasis and promoting longevity (Pellegrino *et al*. 2013; Jovaisaite *et al*. 2014). In contrast, certain neurodevelopmental cell-cell adhesion molecules (e.g. DSCAMs, cadherins, and teneurins) leverage developmental noise to create more robust neural networks. In doing so, they drive repeatable non-genetic phenotypic variation in behavioral responses to serve as a bet hedging strategy (Vogt *et al*. 2008; Ayroles *et al*. 2015; Hiesinger and Hassan 2018; Honegger and DE Bivort 2018).

Genes that modulate a system’s ability to resist developmental noise or a stressor are hypothesized evolutionary targets (Wagner 2008; Vogt 2015; Menezes *et al*. 2018) and subject to natural selection. And yet, these sources of non-genetic phenotypic variation are poorly understood. Previous studies employed a Genome Wide Association Study (GWAS) framework on trait robustness, demonstrating the strategy’s feasibility to identify significant genetic modifiers (Kain *et al*. 2012; Ayroles *et al*. 2015; Morgante *et al*. 2015; Menezes *et al*. 2018; Roman *et al*. 2018). Similarly, we sought to elucidate these genetic factors by studying the robustness of flight performance in a GWAS framework. We turned to the Drosophila Genetics Reference Panel (DGRP) lines, a collection of 205 genetically distinct and inbred lines of *D. melanogaster* that represent a snapshot of natural variation in a wild population (Mackay *et al*. 2012; Huang *et al*. 2014). Using a flight column to assay flies’ ability to react and respond to an abrupt drop (Benzer 1973; Babcock and Ganetzky 2014), we tested 197 DGRP lines for their mean-normalized standard deviation (coefficient of variation) in flight performance. The natural log-transformed coefficient of variation serves as a more normally-distributed proxy for studying phenotypic robustness for genetically distinct groups comprised of genetically identical individuals. In this study, we identified significant individual variants and epistatic interactions, while also exploring the top hits from a whole gene screen across four sex-based phenotypes (males, females, and the average (sex-average) and difference (sex-difference) between sexes). We also used a panel of insertional mutations in several candidate genes (*bru1, CadN, CG15236, CG32181/Adgf-A/Adgf-A2, CG3222, flippy*/*CG9766, CREG, Dscam4, flapper*/*CG11073, Form3, fry, Lasp/CG9692, Pde6, Snoo*), detected in a previous study though they were not significant in the current one (Spierer *et al*. 2021). The successful validation of these genes hints at the dual importance of genes modulating a trait mean and its variance, and it highlights how there are still many more genetic modifiers that affect robustness of flight performance. Across these analyses, we found consistent evidence for the development and function of sensory organs that process external stimuli, including those involved in touch, sight, smell, and sound. Together, these genes highlight the importance of processing proprioceptive cues for robust flight performance.

## METHODS

### Drosophila Stocks and Husbandry

197 Drosophila Genetic Reference Panel (DGRP) lines (Huang *et al*. 2014) and 24 stocks used in the validation experiment were obtained from Bloomington Drosophila Stock Center (Table S1; https://bdsc.indiana.edu/). Flies were grown on a standard cornmeal media (Mossman *et al*. 2016) at 25° under a 12h:12h light-dark cycle. Two to three days post-eclosion, flies were sorted by sex under light CO_2_ anesthesia and given five days to recover before assaying flight performance. All flies scored for robustness, whether in the initial phenotyping screen or in the validation screen, were reared under the same conditions.

### Flight performance assay

We tested approximately 100 flies of each sex from 197 DGRP genotypes (Table S1) using a refined protocol (Babcock and Ganetzky 2014) for measuring flight performance (Benzer 1973). For each sex-genotype combination, groups of 20 flies in five glass vials were knocked down, uncorked, and rapidly inverted down a 25 cm chute. The vials traveled until they reached a stop, at which point flies were ejected into a 100 cm long by 13.5 cm wide tube. Freefalling flies instinctively attempt to right themselves and land. A transparent acrylic sheet coated in TangleTrap^®^ adhesive lined the inside of the tube and immobilized flies at their respective landing height. The sheet, was removed, pinned to a white poster board, and photographed using a Raspberry Pi (model 3 B+) and PiCamera (V2). The positional coordinates were extracted using ImageJ/FIJI’s (Schindelin *et al*. 2012) ‘Find Maxima’ feature with options for a light background and noise tolerance of 30. The distributions of landing heights for each sex-genotype combination were used to calculate the mean distance traveled and standard deviation. The coefficient of variation represents the standard deviation normalized by the mean distance traveled. Finally, we performed a natural log-transformation on each genotype score to make the data more normally distributed. Thus, the natural log-transformed coefficient of variation serves as our phenotype proxy for robustness of flight performance.

### Genome wide association mapping

Robustness phenotypes (Table S2) were submitted to the DGRP2 webserver (reference genome FB5.57) for the association analysis (http://dgrp2.gnets.ncsu.edu/) (Mackay *et al*. 2012; Huang *et al*. 2014). This computational pipeline returned single variant results for four sex-based phenotypes: males, females, average between sexes (sex-average) and difference between sexes (sex-difference). We refer to this analysis and its respective results as the individual variant analysis, since this analysis is the standard when working with the DGRP lines. We analyzed 1,901,174 common variants (minor allele frequency ≥ 0.05) using a mixed effect model to account for Wolbachia infection status and presence of five major inversions. Since certain inversions covaried with the robustness phenotype (Table S4), only significance scores from a linear mixed model accounting for Wolbachia status and the presence of five major inversions were considered.

### Validating candidate genes

Candidate genes (Table S1B) were selected if they were identified from variants identified in the sex-average, individual variant screen for mean landing height and if there were publicly available lines containing a *Minos* enhancer trap (*Mi{ET1}*) mutational insertion (Metaxakis *et al*. 2005) generated by the Drosophila Gene Disruption Project (Bellen *et al*. 2011). Experimental and control lines were derived from common isoparental crosses for each candidate gene stock backcrossed for five generations to the respective w^1118^ or y^1^w^67c23^ background. Isoparental crosses between the resulting heterozygous offspring were partitioned for absence (control line) or presence (experimental line) of the *Mi{ET1}* construct. Experimental lines were verified for homozygosity if all progeny contained the insertion after several rounds of culturing. Validations were conducting in the flight performance assay described above. The distributions in landing heights were assessed for significance if they passed a *P* ≤ 0.05 significance threshold in a Kolmogorov-Smirnov test comparing control and mutant genotypes (Spierer *et al*. 2021).

### Calculating gene-score significance

Gene-level significance scores (gene-score) were determined using `PEGASUS_flies` (Spierer *et al*. 2021), a Drosophila-optimized method for the human-based platform Precise, Efficient Gene Association Score Using SNPs (`PEGASUS`) (Nakka *et al*. 2016). This analysis calculates gene-scores for each gene as a test of whether the distribution of individual variants within a gene (accounting for linkage disequilibrium) deviates from a null chi-squared distribution. Variants from the individual variant association screen were considered and mapped onto gene annotations and linkage disequilibrium files available with the `PEGASUS_flies` package—derived initially from the DGRP2 webserver. Because no variants passed the strict Bonferroni significance threshold (*P* = 3.13E-6), we explored the top five genes for each sex-based phenotype.

### Screening for epistatic interactions

Epistatic hub variants, corresponding with variants more likely to interact with other variants, were identified using MArginal ePIstasis Test (`MAPIT`) (Crawford *et al*. 2017). This approach tests the marginal effect of each variant against a focal phenotype. `MAPIT` requires a complete genotype-phenotype matrix so the DGRP genome was imputed for missing variants using `BEAGLE` 4.1 (Browning and Browning 2007; Browning and Browning 2016) and filtered for MAF ≥ 0.05 using `VCFtools` (v.0.1.16) (Danecek *et al*. 2011).

`MAPIT` was run using the ‘Davies’ method on the raw phenotype scores, 1,952,233 `BEAGLE`-imputed and filtered variants, and the DGRP2 webserver’s relatedness and covariate status files. Since none of the epistatic hub variant *P*-values passed the strict Bonferroni significance threshold (*P* < 2.56e-8) in any of the sex-based phenotypes, we used the 15 most significant variants as a focused subset for targeted pairwise epistasis testing against the unimputed variants (n = 1,901,174). Epistatic interactions were calculated using the ‘–epistasis’ test in a ‘–set-by-all’ framework in `PLINK` (v.1.90) (Purcell *et al*. 2007). Significant epistatic interactions were considered if they passed a Bonferroni threshold (*P* < 1.75E-9).

### Data availability

All phenotype data required to run the outlined analyses are available in Table S2 or using the DGRP2 webserver (http://dgrp2.gnets.ncsu.edu/).

## RESULTS and DISCUSSION

We sought to identify the genetic modifiers of robustness in a whole organism phenotype: flight. Using the Drosophila Genetic Reference Panel (DGRP) lines, we identified several individual variants, validated a previously identified subset of genes for robustness, and two pairs of significant epistatic interactions. While we didn’t find any significant whole genes, some of the most significant genes corresponded with modifiers of trans-regulatory gene expression and detecting external stimuli. In the sections that follow we describe the variant-based analysis, gene-based analysis, epistatic analysis, and validation of candidate genes.

### Variation in flight performance across the DGRP

We screened 197 DGRP lines (Table S1A) for their flight ability in response to an abrupt drop (Figure 1A-B). Qualitative observations made in a previous study of strong, intermediate, and weak genotypes in the flight assay suggests stronger genotypes react faster and respond more effectively than weaker one (Spierer *et al*. 2021). The mean and standard deviation in landing height were calculated for each sex-genotype combination, though the standard deviation was related to the mean landing height (males: R = 0.72, *P* < 1E-32; females: R = 0.68, *P* < 1E-28). To study variation in the absence of the mean, we chose to use the coefficient of variation. Additionally, we natural log-transformed the coefficient of variation to make the data more normally distributed (Figure S1). An earlier pre-print of this work calculated the coefficient of variation as the standard deviation normalized by the mean landing height from the bottom of the column (Spierer and Rand 2020), though this created a negative association between our metric and robustness so we chose to normalize the standard deviation by the mean distance fallen in the column (Table S2). Thus, the natural log-transformed coefficient of variation served as our metric for robustness.

**Figure 1.**
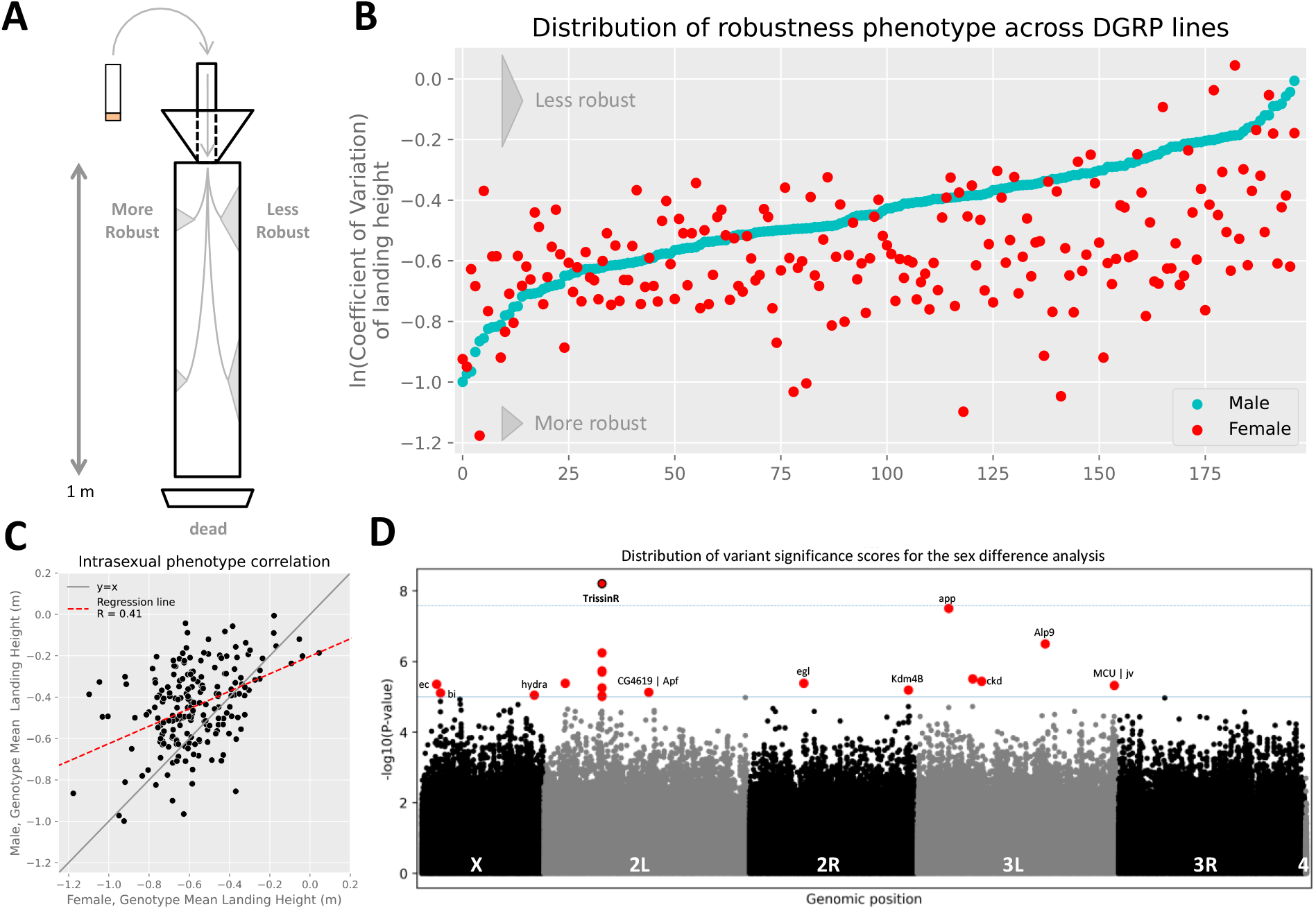
The *Drosophila* Genetic Reference Panel lines demonstrate variation for robustness in flight performance across genotypes and sexes. (A) Flies were assayed for flight performance using a meter-long flight column (Babcock and Ganetzky 2014). The natural log-transformed coefficient of variation (mean-normalized standard deviation) is a proxy for robustness; more robust genotypes have less variation in landing height around the mean. Flies that passed through the column were excluded from the analysis. (B) The phenotypic distribution of sex-genotype pairs, ordered by increasing male score, demonstrates the DGRP lines have variation in their robustness for flight performance. Genotypes demonstrated phenotypic variation for robustness in both sexes. More negative values correspond with increased robustness. (C) Males were generally more robust than females, though the two were related (R = 0.41, *P* < 5E-9; regression line in red). Sexual dimorphism is observed by the intersection of the regression line and y = x line (gray). (D) Individual variants in the sex-difference analysis, visualized as a function of the –log10 of variants’ *P*-value illustrates several variants (red) passed the suggestive DGRP significance threshold (*P* ≤ 1E-5; blue solid line), and one (red with black outline) passed Bonferroni significance threshold (*P* ≤ 2.63E-8, blue dashed line). Variants that did not pass the significance threshold are colored in black or gray by chromosome. Other sex-based phenotype Manhattan plots are available in Figure S4.

In this study, genotypes with a lower coefficient of variation (more consistent) were more robust for flight performance (Klingenberg 2019). On average, flight performance was more robust in males than females (males: -0.45 A.U. ± 0.19 SD vs. females: -0.58 A.U. ± 0.19 SD; Figures 1B). There was a significant relationship in robustness between sexes (R = 0.41; *P* < 5E-9; Figure 1C), suggesting the genetic architecture of robustness in flight performance is similar between the sexes. However, the magnitude of the regression coefficient suggests robustness is somewhat sexually dimorphic.

We tested our phenotype in both males and females against those publicly available on the DGRP2 webserver to determine whether robustness of flight performance was a unique trait. We found no significant relationship after imposing a significance threshold of *P* ≤ 1.8E-3 to account for multiple testing (Table S3), suggesting our phenotype is unique.

### Several variants of large effect associate with robustness in flight performance

We performed a Genome Wide Association Study (GWAS) to calculate each variant’s significance, and subsequently whole gene significance scores. We analyzed the effects of 1,901,174 common variants (MAF ≥ 0.05) across for four sex-based phenotypes (males, females, the sex-average, and sex-difference; Figures 1D and S2-4). Although none of the major inversions or presence of Wolbachia covaried with our phenotype scores (Table S4), we still used a mixed effects model to minimize extraneous sources of variation.

We performed a GWAS on the robustness phenotype using the DGRP2 webserver pipeline. Only one variant (2L_5852054_SNP; *P* = 6.24E-9) in the sex-difference screen passed a strict Bonferroni significance threshold (P = 2.63E-8). This variant mapped to an intron of *TrissinR*, a neuropeptide receptor that binds *Trissin* and acts as a G-protein coupled receptor. *TrissinR* was previously identified to be important for neuronal communication in Olfactory Receptor Neurons (ORN) and Ionotropic Receptors (IR) (Mclaughlin *et al*. 2021). The gene’s importance in flight was previously documented in our previous study on the genetic modifiers of the mean of flight performance where it was a significant epistatic interactor with the chemo- and mechanosensing gene *ppk23* (Spierer *et al*. 2021).

Applying the individual variant, DGRP association threshold (*P* ≤ 1E-5), we identified 69 unique variants across 41 genes (Table S5). No variant corresponded with protein coding changes, suggesting variation in complex traits is driven by modulation of gene regulation rather than changes to protein coding sequence (Mackay *et al*. 2012; Mackay and Huang 2018). Seventeen of these genes were identified from several different analyses in our prior study: *app, CG10362, CG15270, CG17839, CG32264, CG43313, cv-c, dpr2, ec, Eip75B, Gmap, jv, Kdm4B, ncd, ppk8, TrissinR, X11Lbeta*. This overlap is suggestive that genes affecting a trait mean may also be important for affecting variation in the same trait.

In addition to direct overlaps in genes, we identified four paralogous genes shared between the present and prior study. In the present study, *Dscam2* is a paralog with *Dscam4*, which was identified in the individual variant analysis as a Bonferroni variant and validated for its role in mean flight performance. Dscam genes are also paralogs with defective proboscis response (dpr) genes, like *dpr2*, which was also identified here. Finally, two pickpocket genes (*ppk8* and *ppk27*) were paralogous with *ppk23*, a highly significant epistatic hub gene that is likely involved in relaying proprioceptive information.

### Analyses of whole-gene effects identifies distinct factors affecting robustness

The individual variant screen takes a minSNP approach, deeming a gene significant if its most significant variant passes a significance threshold. However, this approach is biased toward longer genes and does not account for linkage between sites. To counteract these biases, we employed `PEGASUS_flies` (Spierer *et al*. 2021), a *Drosophila* version of the human-focused `PEGASUS` platform (Nakka *et al*. 2016). This method takes a gene-specific approach; assessing a whole gene’s significance by testing the distribution of variants within a gene against a null chi-squared distribution of SNP *P*-values. Thus, it can detect significant genes of moderate effect, as well as genes that may be missed in a minSNP approach.

We failed to identify any significant genes across the four sex-based phenotypes using a strict Bonferroni threshold (*P* ≤ 3.43E-6). Since this threshold is overly conservative, we looked at the top five genes from each of the four sex-based analyses and identified 18 unique genes using `PEGASUS_flies` (Table S6 and Figure S5). Of these genes, only one had a single *P*-value exceed the individual significance threshold (*P* = 1E-5; *jv* in sex-average), demonstrating how `PEGASUS_flies` is capable of expanding the list of potential candidate genes in GWAS-type studies.

Of the top five genes in each sex-based phenotype, four corresponded with trans-regulatory factors (*CG2034, CG4565, CG42526, Wdr82*) (Gaudet *et al*. 2011), which supports our earlier observation that variants identified through the individual variant analysis were in non-coding regions. Additionally, we identified genes involved in sensing the external environment through the chaeta development (hair-like structures responsible for chemo- and mechanosensation; *jv*) and the development of chordontal organs (stretch receptor organs; *btv*) (Eberl *et al*. 1997; Eberl *et al*. 2000; Shapira *et al*. 2011). While these genes were not significant under a strict Bonferroni significance threshold, they still support an important role for variation in proprioception and receiving external stimuli in modulating the robustness of flight performance.

### Association of epistatic hub and pairwise epistatic variants with robustness in flight performance

Epistatic, or pairwise, interactions play an outsized role as context-specific effectors in complex traits (Huang *et al*. 2012). Traditional epistasis analyses face large computational and statistical hurdles, so we turned to MArginal ePIstasis Test (`MAPIT`) to focus the exhaustive pairwise search and identify epistatic hub variants with a greater likelihood of interacting with other variants (Crawford *et al*. 2017). These hub variants were then used as a subset in a set-by-all pairwise epistasis search against all variants considered in the individual variant association analysis.

We failed to identify any epistatic hub variants that passed a strict Bonferroni significance threshold (*P* = 2.56E-8). Since we were using `MAPIT` to narrow our search space for epistatic variants, we decided to focus on the 15 most significant variants in each sex-based phenotype to inform our search for epistatic variants instead. Doing so yielded two pairs of epistatic interactions, one in each the female and sex-difference analyses though none leading to changes in the protein coding sequence (Table S7).

The female interaction was between SNP pairs X_14165625_SNP and 2R_3523428_SNP. The former corresponded with a synonymous coding site in *narrow abdomen* (*na*), an ion channel involved in locomotor rhythm and mechanosensation (Nash *et al*. 2002; Lear *et al*. 2013). The latter corresponded with two separate genes, Myosin-7a binding protein (*M7BP*; intron) and *antisense RNA:CR45131* (704 bp upstream). Interestingly, *M7BP* localizes to actin-bundles in sensory organs in *Drosophila*, as well as the Johnston’s organ, which is used for auditory sensation (Kiehart *et al*. 2004; Todi *et al*. 2005; Todi *et al*. 2008; Liu *et al*. 2021). The connection between *na* and *M7BP* supports the importance of sensory hairs in proprioception and receiving external stimuli during flight that may modulate robustness.

The sex-difference interaction was between SNP pairs 3L_7643140_SNP and 3R_16731290_SNP. The first SNP lies 508 bp upstream of *CG32373*. It is expressed in the Johnston’s organ and is hypothesized to aid in synaptic formation (Kurusu *et al*. 2008; Senthilan *et al*. 2012). Additionally, it is hypothesized to work with *nmo*, previously identified in flight performance (Spierer *et al*. 2021), in ommatidial rotation (MUNOZ-Soriano *et al*. 2013). Meanwhile, 3R_16731290_SNP falls within or near two genes: *Turandot X* (*TotX*) and *Grik. TotX* is a stress response gene in the JAK-STAT pathway best known in the context of heat stress (Manenti *et al*. 2018), though it has been documented to have some connection to auditory processing (Immonen and Ritchie 2012). While it is possible that it interacts with *CG32373*, it is far more likely that the main epistatic interaction is with *Grik*, a glutamate receptor involved in synaptic transmission in the adult brain and visual system (Gaudet *et al*. 2011; Karuppudurai *et al*. 2014). It is orthologous to *glutamate ionotropic receptor kainate type subunits 1-3* (*GRIK1-3*), involved in the development of intellectual disability and Huntington’s disease in humans (Macdonald *et al*. 1999). Together, it would follow that *CG32373* and *Grik* might work together in the *Drosophila* flight system to process visual and/or auditory signals that are important in the robustness of flight performance.

### Functional validation of candidate genes supports a role for neurodevelopment affecting robustness of flight performance

Finally, we sought to test whether genes that modify the mean flight performance phenotype also modify the robustness in flight performance. To do so, we tested 24 independent insertional mutations in candidate genes identified from an earlier study on mean flight performance (Spierer *et al*. 2021). Of these, 21 constructs fell in unique genes while three constructs were used as independent replicates of different highly significant genes in the mean flight phenotype (*CadN, Dscam4, flap* (*CG11073*)) (Spierer *et al*. 2021). Of the 21 unique genes, all but one (*CREG*) were strongly significant in the mean flight performance paper’s list of top variants (Spierer *et al*. 2021). Despite their significance in the other study, none of these genes were significantly associated with robustness in any of the four sex-based phenotypes in the present study. Thus, we were also able to test whether there were significant genes affecting robustness that we were unable to detect due to a lack of power.

Of these 21 genes, there was a significant difference in robustness for 13 constructs using a comparison of genotypes carrying an insertional mutation in a candidate gene of interest against their backcross-control genotypes. We found statistical significance with a Kolmogrov-Smirnov test for 11 candidate genes where the construct inserted within single genes (*bru1, CadN, flip* (*CG9766*), *CG15236, CREG, Dscam4, flap* (*CG11073*), *form3, fry, Pde6*, and *Snoo*) and two where the construct inserted in multiple genes (*Adgf-A/Adgf-A2/CG32181 and CG9692/Lasp*) (Figure 2 ; Table S8). These genes were also previously validated in the mean flight performance screen (Spierer *et al*. 2021), suggesting that genes likely play dual roles modifying the ability and variability of flight performance. These analyses using insertional mutations showed that while natural variation in this set of 21 candidate genes for mean flight performance do not pass robustness of flight GWA thresholds for significance, specific mutations in those genes are capable of impacting robustness in 13 of these 21 genes.

**Figure 2.**
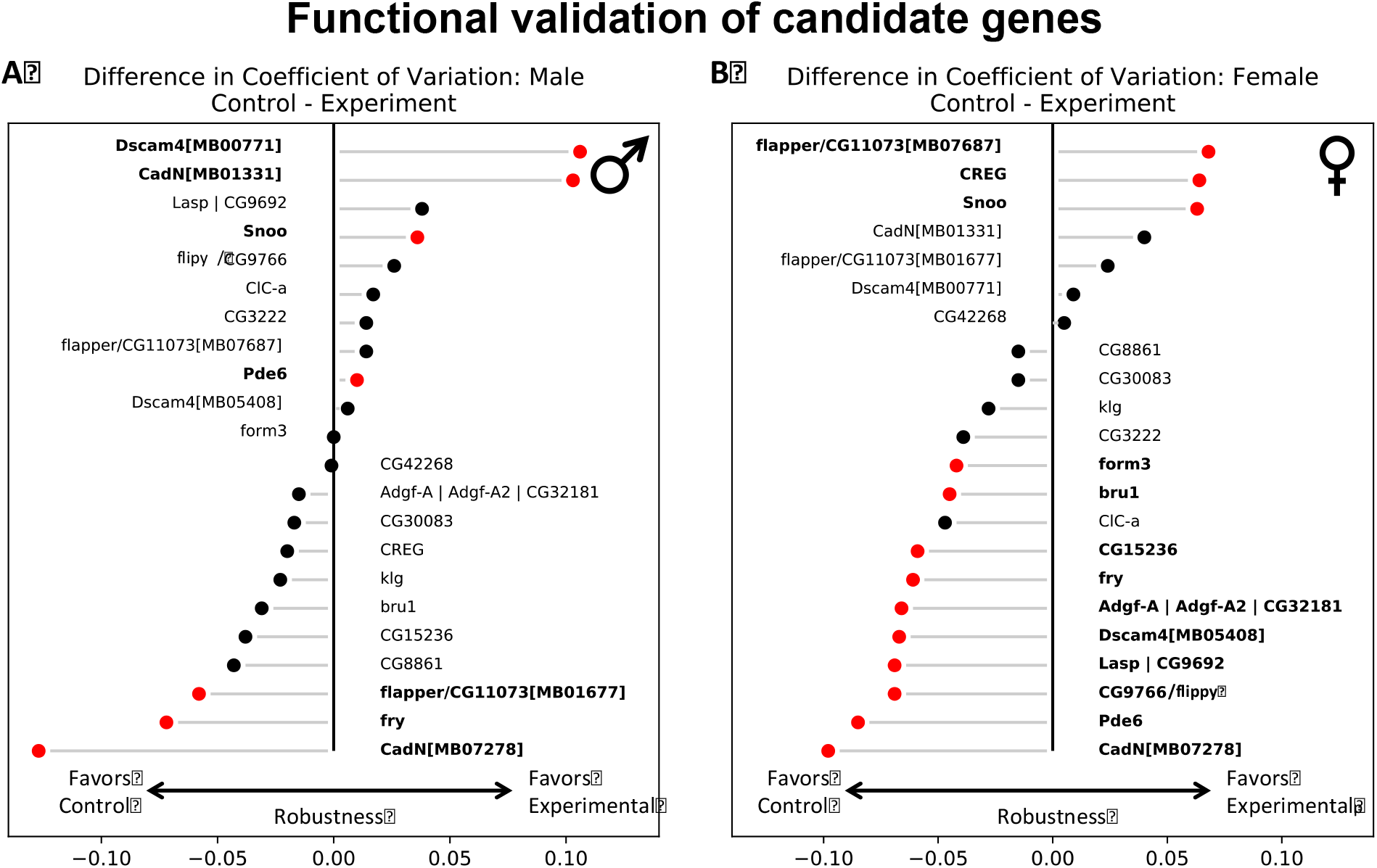
Several genes validated for robustness of flight performance. Flies homozygous for *Mi{ET1}* insertion constructs inserted in candidate genes (experiment) were tested against their background control (control). Comparisons between control and experiment lines were assessed for significance using a Kolmogrov-Smirnoff test (*P* ≤ 0.05; red points and bold text). Values to the left of the midline suggest control genotypes were more robust than experimental lines, while the opposite is true for values to the right of the line. (A) Seven constructs were significant in males, (B) while 13 were significant in females. Some candidate genes were tested more than once (*CadN, Dscam4*, and *flap*) because they were strongly significant in the sex-average, individual variant association screen. Separate constructs are denoted by a suffix containing a ‘MB’ code.

Interestingly, *CadN* and *Dscam4* are important genes contributing to type IV dendritic arborization sensory neurons. These genes are known to contribute to robustness as they connect sensory structures (e.g. chaete) to the peripheral nervous system. They also work with teneurins (e.g. *Ten-a*), which are known to affect robustness of locomotor handedness (Buchanan *et al*. 2015). *CadN* and *Dscam4* also work with *fry* and *Snoo*, which develop and pattern chaete and campaniform sensilla on the wing, and are likely useful in mechanosensation and proprioception during flight (Emoto *et al*. 2004; Neves *et al*. 2004; Soba *et al*. 2007; Fuerst and Burgess 2009; Quijano *et al*. 2010; Matsubara *et al*. 2011).

Our findings suggest that our experimental design is sufficient to identify individual variants affecting robustness. However, we were limited in our power to detect whole gene or epistatic interactions affecting robustness. While we could not comprehensively detect all genetic modifiers of flight robustness, the fact that mutations in genes affecting mean flight performance can affect robustness implies that many other genes affecting robustness likely exist. Even using all but eight of the available DGRP lines, we lacked the power to detect many genes. Therefore, we suggest that future studies exploring the mean and robustness for traits with the DGRP lines should supplement the core panel with other sources of genetic variation, such as the Global Diversity Panel (GDP) or an Advanced Intercross Population (AIP) (Grenier *et al*. 2015; Mackay and Huang 2018).

### Conclusions

We present results from four analyses across four sex-based phenotypes surveying different facets of the genetic architecture of robustness for flight performance. The individual variant analysis was the most fruitful for identifying novel genetic modifiers of robustness in flight performance, while the screen for epistatic interactions found two pairs of genes that were both involved in processing external cues (mechano-, audio- and visuosensory sensory) that are also likely important for proprioception. A whole gene screen did not meet strict significance thresholds though the most significant genes in the analysis indicated trans-regulatory genes and some genes involved in the development of proprioceptive structures were important. Finally, we validated several genes roles in contributing to robustness of flight performance that were not detected in this study. This result suggests that despite our current findings, there are many more genetic modifiers of robustness left to identify. These genetic modifiers likely require additional genotypic and phenotypic variation to detect, so we suggest future studies supplement the DGRP with other panels of flies (GDP or an AIP) to counteract these limitations.

Future studies in other phenotypes should consider evaluating both the mean and standard deviation or coefficient of variation for their focal phenotype to better understand modifiers affecting robustness in a specific complex trait, as well as robustness in complex traits more generally. Doing so would provide a better survey of the genetic modifiers of robustness as a phenotype and allow for greater insights into the mechanisms of evolutionary change.

## Author contributions

DMR and ANS conceived the idea and designed the experiment. ANS performed experiments and analyses. DMR and ANS wrote and revised the manuscript.

## Data accessibility statement

All phenotype data required to run the outlined analyses are available in Table S1 or using the DGRP2 webserver (http://dgrp2.gnets.ncsu.edu/). Supplemental tables and supplemental figures are hosted by Dataverse: https://doi.org/10.7910/DVN/MV7QA4.

## Acknowledgements

We thank Faye Lemieux for her assistance with fly husbandry and Jim A. Mossman for his assistance collecting phenotype data.

